# The essential genome of *Escherichia coli* K-12

**DOI:** 10.1101/237842

**Authors:** Emily C. A. Goodall, Ashley Robinson, Iain G. Johnston, Sara Jabbari, Keith A. Turner, Adam. F. Cunningham, Peter A. Lund, Jeffrey A. Cole, Ian R. Henderson

**Author notes:** Address correspondence to Ian R. Henderson. E.C.A.G. and A.R. contributed equally to this work.

## Abstract

Transposon-Directed Insertion-site Sequencing (TraDIS) is a high-throughput method coupling transposon mutagenesis with short-fragment DNA sequencing. It is commonly used to identify essential genes. Single gene deletion libraries are considered the gold standard for identifying essential genes. Currently, the TraDIS method has not been benchmarked against such libraries and therefore it remains unclear whether the two methodologies are comparable. To address this, a high density transposon library was constructed in *Escherichia coli* K-12. Essential genes predicted from sequencing of this library were compared to existing essential gene databases. To decrease false positive identification of essential gene candidates, statistical data analysis included corrections for both gene length and genome length. Through this analysis new essential genes and genes previously incorrectly designated as essential were identified. We show that manual analysis of TraDIS data reveals novel features that would not have been detected by statistical analysis alone. Examples include short essential regions within genes, orientation-dependent effects and fine resolution identification of genome and protein features. Recognition of these insertion profiles in transposon mutagenesis datasets will assist genome annotation of less well characterized genomes and provides new insights into bacterial physiology and biochemistry.

**IMPORTANCE:** Incentives to define lists of genes that are essential for bacterial survival include the identification of potential targets for antibacterial drug development, genes required for rapid growth for exploitation in biotechnology, and discovery of new biochemical pathways. To identify essential genes in *E. coli*, we constructed a very high density transposon mutant library. Initial automated analysis of the resulting data revealed many discrepancies when compared to the literature. We now report more extensive statistical analysis supported by both literature searches and detailed inspection of high density TraDIS sequencing data for each putative essential gene for the model laboratory organism, *Escherichia coli*. This paper is important because it provides a better understanding of the essential genes of *E. coli*, reveals the limitations of relying on automated analysis alone and a provides new standard for the analysis of TraDIS data.

## INTRODUCTION

There are many incentives to define lists of genes that are either essential for bacterial survival or important for normal rates of growth. Essential genes of bacterial pathogens may encode novel biochemical pathways or potential targets for antibacterial drug development. Disruption of genes required for rapid growth results in strains handicapped for exploitation in biotechnology. Conversely, normal growth of mutants defective in genes previously expected to be essential could reveal unexpected parallel biochemical pathways for fulfilling the essential function.

Multiple attempts have been made to generate definitive lists of essential genes, but there are still many discrepancies between studies even for the model bacterium *Escherichia coli* strain K-12. Two general approaches have been used: targeted deletion of individual genes, as in the Keio collection of mutants (1); and random mutagenesis (2, 3). Data from several studies using different mutagenesis strategies have yielded inconsistent data and hence conflicting conclusions. <u>T</u>ransposon <u>D</u>irected <u>I</u>nsertion-site <u>S</u>equencing (TraDIS) is one of several high-throughput techniques that combine random transposon mutagenesis with sequencing of the transposon junctions in high density mutant libraries (4–7). Since its inception in 2009, this high-throughput method has been applied to a range of biological questions (4, 8–15). Here, in order to resolve outstanding conflicts, we report the use of this approach to identify the essential genes of *Escherichia coli* K-12 strain BW25113, a well-studied model organism for which a complete gene deletion library is available (1).

A confounding factor in determining the ‘essentiality’ of a gene is the definition of an essential gene. Complete deletion of an essential gene results, by definition, in a strain that cannot be isolated following growth. However, it is well known that certain genes are required for growth under specific environmental and nutritional conditions. Such genes can be considered conditionally essential. For the purposes of this study we define a gene as essential if the transposon insertion data reveal that the protein coding sequence (CDS), or a portion of the CDS, is required for growth under the conditions tested here. To aid our analysis we developed a statistical model that included corrections for both gene length and genome length in order to decrease false positive identification of essential genes.

An additional challenge with defining essentiality in high throughput studies is an over-reliance on automated analysis of the data. For example, a consequence of relying only on quantification of the number of unique insertions within a gene is that genes with essential regions will be missed. If only part of a gene encodes the essential function, it should be possible to isolate viable mutants with transposon insertions in non-essential regions of the coding sequence (2). Conversely, reliance on statistical analysis alone can also lead to over-estimation of the number of essential genes. This is a common result from insertion sequencing analysis (16). A low number of transposon insertion events within a gene, which fall below the statistical cut-off threshold, can be due to inaccessibility of the gene to transposition because of extreme DNA structure, exclusion by DNA-binding proteins, polarity effects due to insertion in a gene upstream of a co-transcribed essential gene, and location of the gene close to the replication terminus (17). The most frequent reason for a low number of insertions is that the product of the disrupted gene is required for normal rates of growth under the conditions tested. In the current study, to minimize the possibility of incorrectly designating genes as essential or contributing to fitness, we have supported our statistical analysis with a gene-by-gene inspection of the insertion distribution within each individual gene.

## RESULTS AND DISCUSSION

### Sequencing of a mini-Tn*5* transposon insertion library in *E. coli* strain BW25113

We have used a modified method to obtain TraDIS data for a transposon mutant library of *E. coli* K-12 strain BW25113 (4, 9). The strain BW25113 was chosen because it is the parent strain for the Keio collection of deletion mutants and ideal for a direct comparison between datasets. A mini-Tn*5* transposon with a chloramphenicol resistance cassette was transformed into competent cells and grown overnight on selective medium. Individual colonies were pooled to construct the initial library, estimated to consist of approximately 3.7 million mutants. An Illumina MiSeq was used to obtain TraDIS data from two independent DNA extracts of the transposon library, NTL1 and NTL2 (Table 1). Raw data were checked for the presence of an inline index barcode to identify independently processed samples (Table 1). This resulted in 4,818,864 sequence reads from NTL1 and 6,189,409 from NTL2. After verification of the presence of a transposon sequence and removal of poor quality data or short sequence reads, 3,891,339 (80.75%) and 4,387,970 (70.89%) sequence reads, respectively, were mapped successfully to the *E. coli* K-12 BW25113 genome (CP009273.1; Table 1). The distribution of insertion sites covers the full length of the genome (Fig. 1A). There was a high correlation coefficient of 0.96 between the samples (Fig. 1B). The data were therefore combined to give a total of 8,279,317 sequences that were mapped to 901,383 unique insertion sites throughout the genome. Similar numbers of insertions, 481,360 and 480,072, were found for both orientations of the transposon. The high density of unique insertion sites resulted in an average of 1 insertion every 5.14 bp and a median distance between insertions of 3 bp. An example is shown in Fig. 1D.

**Figure 1.**
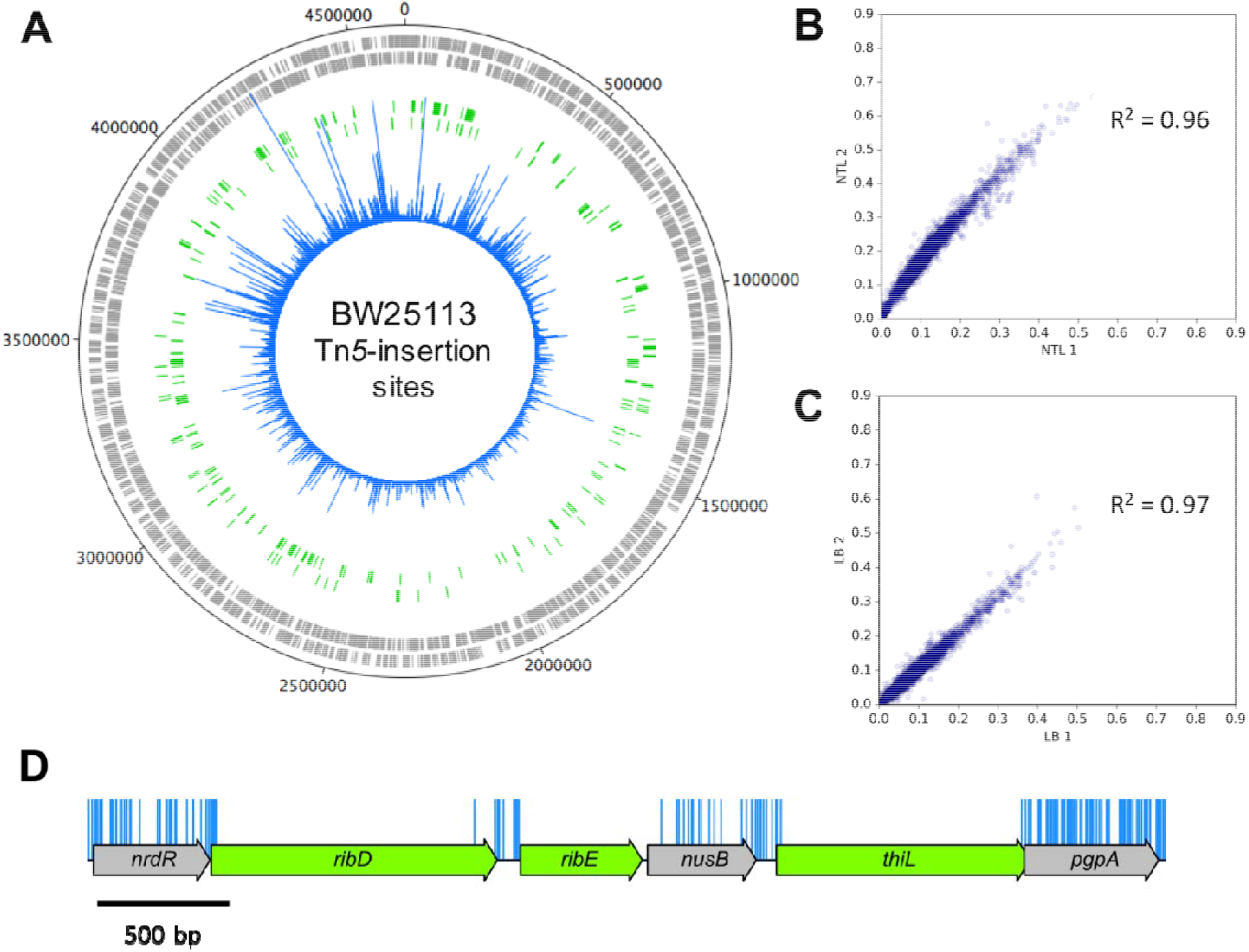
Genome wide transposon insertion sites mapped to *E. coli* strain BW25113. (A) Frequency and location of transposon junction sequences from a mini-Tn*5* transposon library in strain BW25113, mapped to the BW25113 genome (CP009273.1). The outer track marks the BW25113 genome in bp starting at the annotation origin. The next two inner tracks correspond to sense and anti-sense CDS respectively (grey), followed by two inner tracks depicting the essential genes identified by TraDIS on the sense and anti-sense strands, respectively (green). The innermost circle (blue) corresponds to the frequency and location of transposon insertion sequences mapped successfully to the BW25113 genome after identification of a transposon sequence. Figure created using DNAPlotter. (B) The correlation coefficient of gene insertion index scores for two sequenced technical replicates of the input transposon library (NTL1 and NTL2) (C) and following growth in LB (LB1 and LB2). (D) Representation of transposon insertion points across a portion of the K-12 BW25113 genome (blue), showing essential genes (green) and non-essential genes (grey). Blue bars correspond with transposon insertion sites along the genome and have been capped at a frequency of 1.

**Table 1.** Parameters for TraDIS data set derived from the *E. coli* K-12 strain BW25113 transposon library

| Condition | Number of sequence reads with matching inline barcode | Mapped sequence reads | Number of genome-wide insertion sites |
| --- | --- | --- | --- |
| NTL1 | 4,818,864 | 3,891,339<br>(80.75% of the raw data) | 500,476 |
| NTL2 | 6,189,409 | 4,387,970<br>(70.89% of the raw data) | 817,011 |
| NTL combined | 11,008,273 | 8,279,309 | 901,383 |
| LB1 | 5,908,163 | 5,201,711<br>(88.04% of the raw data) | 400,009 |
| LB2 | 6,403,324 | 5,382,477<br>(84.06% of the raw data) | 419,660 |
| LB combined | 12,311,487 | 10,584,188 | 595,233 |

### Identification of putative essential genes by TraDIS

To determine whether a gene was essential or non-essential the numbers of insertions per CDS were quantified. CDS is defined as the protein coding sequence of a gene, inclusive of the start and stop codons. To normalize for gene length the number of unique insertion points within the CDS was divided by the CDS length in bases. This value was termed the insertion index score and has been used previously as a measure of essentiality (4, 8, 9, 18), given a sufficiently dense library (19).

The frequency distribution of the insertion index scores was bimodal (Fig. S1), as previously shown by others (2). We assume that genes in the left-hand mode, which have a low number of transposon insertions, are either essential for survival or genes that, when disrupted, confer a very severe fitness cost (Fig. 1D). The second mode contains genes with considerably more insertions; these are deemed non-essential (Fig. 1D). Based on inspection of the distributions, an exponential distribution model was fitted to the mode that includes essential genes and a gamma distribution model was fitted to the non-essential mode. For a given insertion index score, the probability of belonging to each mode was calculated and the ratio of these values was termed the log-likelihood score. A gene was classified as essential if its log-likelihood score was less than log_2_(12) and was therefore 12 times more likely to belong to the essential mode than the non-essential mode. Using this approach, sufficient insertions were found in 3793 genes for them to be classed as non-essential, 162 genes were situated between the two modes and classed as ‘unclear’ and 358 genes in the mutant library were identified as essential (Table S1).

The 358 candidate essential genes identified in the NTL data were compared to the essential genes as defined by the Keio collection and the Profiling of the *E. coli* Chromosome (PEC) database (1, 2). This comparison revealed 248 genes (59.5%) that were common to all three datasets (Fig. 2A; Table S2). This agreement between all three datasets strongly supports the hypothesis that these genes are essential so they were not investigated further. An additional 169 genes were identified as potentially essential in only one or two of the datasets. These comprise 16 genes in the Keio and PEC lists that were not identified by our analysis, 25 exclusive to Keio, 18 exclusive to PEC, and 11 and 18 that overlapped between our method and Keio or PEC, respectively (Fig. 2A). However, the largest subcategory of 81 genes is unique to our dataset.

**Figure 2.**
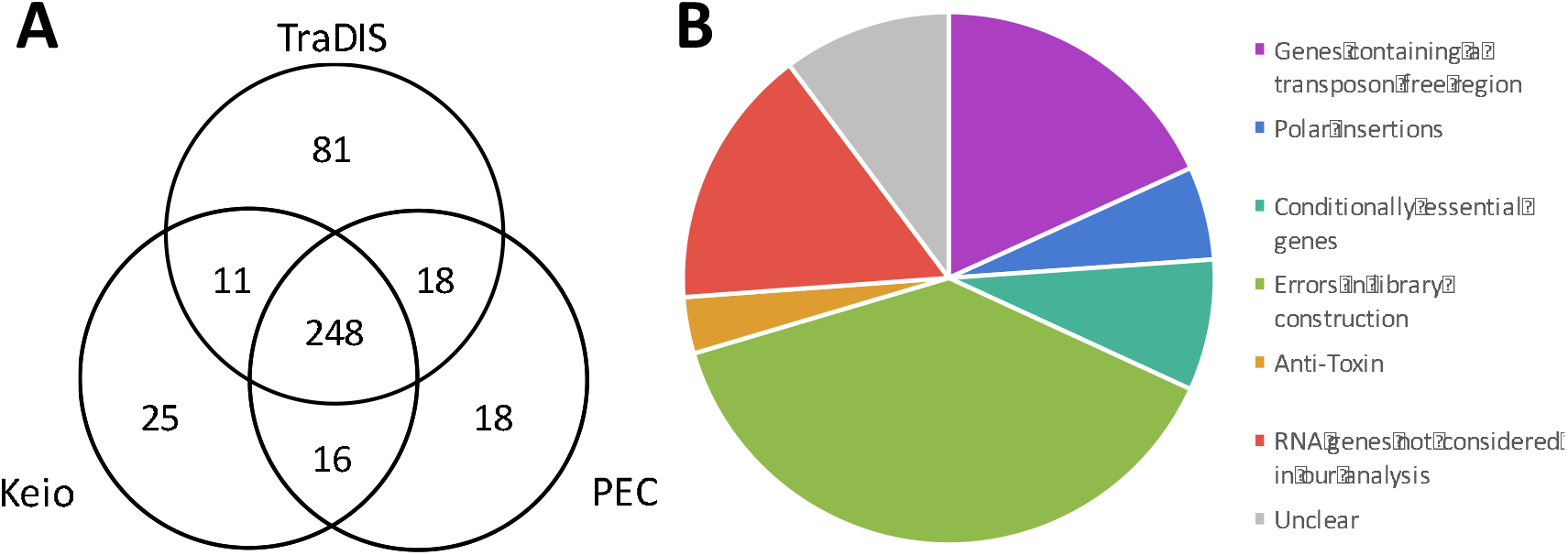
Comparison of essential gene data from various sources and examples of insertion profiles overlooked by automated statistical analysis of insertion index scores. (A) Essential gene candidates identified using TraDIS were compared to existing essential gene data. A 3-way comparison between the Keio collection of single gene knockouts, the Profiling of the *E. coli* Chromosome (PEC) online database and our transposon-insertion sequencing data identified 248 essential genes that were common to all three data. (B) The outlying genes of the Venn diagram, excluding those unique to our TraDIS dataset, were inspected to understand the source of discrepancy between datasets. Genes were grouped into the overarching categories of “Transposon free region” (blue), “Anti-toxin” (green), “Polar insertions” (purple), “Conditionally essential” (orange), and “Errors in library construction” (red) with subcategories depicted in different shades of the colour. Genes not included in our analysis or that remain unclear are shown in yellow or grey respectively.

### Statistical analysis of the transposon insertion density data

Over-estimation of the number of genes that are essential has been noted in studies using transposon insertion sequencing (16). In previous attempts to use statistical analysis to define an essential gene, a Poissonian model was used to derive a p-value for an insertion-free region (hereafter IFR) of a given length against the null hypothesis that, by chance, no insertions occurred in that region. We refined this approach for two reasons. First, genomes are sequences of discrete sites: although a continuous Poisson model can provide an approximation to this structure, a naturally discrete picture is more representative of true genome structure. Second, unless corrections are applied for gene length or for the genome length, this method risks overestimating the total number of essential genes. This problem arises because the method implicitly considers only a single, small genomic region, giving the probability that no insertions will be found in a single region of a given bp length. However, genes and genomes have many such regions that are effectively independent, so the genome-wide probability of observing a “false positive” insertion-free region across the genome will be much higher.

To avoid this risk of over-interpretation of TraDIS data, we propose a new statistical approach, summarized in Supplementary Methods and Supplementary Fig. S2. First, we replaced the commonly-used Poissonian model exp(-f/x) with a geometric model (see (20) for further discussion of this), giving the probability of seeing k-1 "failures" (an insertion-free site) then a “success” (insertion event) in a string of independent trials as P(k) = p (1-p)^(k-1), where p is the probability of a success (here, an insertion). The p-value associated with a string of l sites being insertion-free is then p = sum_(k=l)^(inf) P(k), an easily computable quantity. Next, to guard against false positives, we need to precisely state the statistic of interest and the corresponding null model. Under a null model of random, independent insertions, the three probabilities most pertinent here are those with which (i) a single length-l region has no insertions; (ii) a gene of length g contains one or more insertion-free regions of length l; (iii) a genome of length G contains one or more insertion-free regions of length l. We used stochastic simulations of random insertions with given densities and genome lengths (see Supplementary Methods) to compute these probabilities. These values then give p-values for insertion-free region observations, correcting for gene and genome length. Specifically, p_gene(l) is the probability of observing one or more insertion-free regions of at least length l in a model gene (of length g = 1000bp) by chance (ii), and p_genome(l) is the probability of observing one or more insertion-free regions of at least length l in a full genome (of length G = 4.6Mb) by chance (iii). The uncorrected p-value (i) is that typically reported in other studies. Statistical analysis of our current data (901,383 inserts in a 4,631,469 bp genome) gives a corrected p_genome = 0.05 for l ~= 75 bp and p_gene = 0.05 for l ~= 36 bp (p_gene = 0.005 for l ~= 47 bp). In other words, there is a probability of 0.05 that any insertion-free region of length 75 could appear anywhere in the genome by chance, and there is a probability of 0.005 that any insertion-free region of length 47 will occur anywhere in a gene of length 1000 by chance. To our knowledge, this represents the first study with a confident and genome-wide corrected detection resolution (Fig. S2), and the closest yet to approaching the length of the smallest annotated gene in our reference genome (CP009273.1), which is 45 bp.

In checking for uniformity of insertion density across genomic regions, we found that the density of insertions around the terminus (taken as a region centered around *terABCD*) was slightly lower than the genomic average (a density of 0.142 in the surrounding 500 kb region, or 0.145 in the surrounding 1 Mb region, compared to a 0.195 average; Fig. 1A). This density change marginally increases the detection of false-positive essential genes in the vicinity of the terminus, but still represents an unprecedented level of coverage.

### Resolution of conflicts between datasets

A critical requirement for the validation of a list of essential genes is to explain why the statistical analysis of transposon insertion data failed to identify genes that the Keio library of deletion mutants and the PEC database identified as essential. We coupled statistical analysis and manual inspection of the data with literature searches to rationalize conflicting results. We find that many of the inconsistencies between datasets can be explained by different methodologies used, definitions of the term ‘essential’ and statistical approaches (Fig. 2B).

#### Genes containing transposon free regions

Manual inspection of the data revealed genes with transposon free regions that were large enough to be identified as significant using the algorithm defined in the previous section. These IFRs do not necessarily report that a gene is essential, but that the insertions within these genes are sufficiently sparse that the IFR is unlikely to have occurred by chance. These genes fall loosely into two groups. The first group contains genes for which the 5′ regions are essential and contain no insertions. However, there are transposon insertions in the non-essential regions of these genes, such as *ftsK* (Fig. 3A, Table S3). FtsK is involved in correct segregation of the chromosome during division (20, 21); the N-terminal domain of FtsK contains 4 transmembrane passes and is required for localization of FtsK to the septum (22–24). There is substantial literature reporting the essential function of the N-terminal domain, consistent with our data (21, 22, 25). This is a common observation within insertion data and arises when only the function of the N-terminus of the protein is required for viability (8). Initial analysis of transposon insertion data would lead to these genes being incorrectly classified as non-essential, but attempts to construct a deletion mutant would fail. Indeed, previous transposon sequencing experiments failed to identify the essential nature of some of these genes when relying on statistical analysis alone (9).

**Figure 3.**
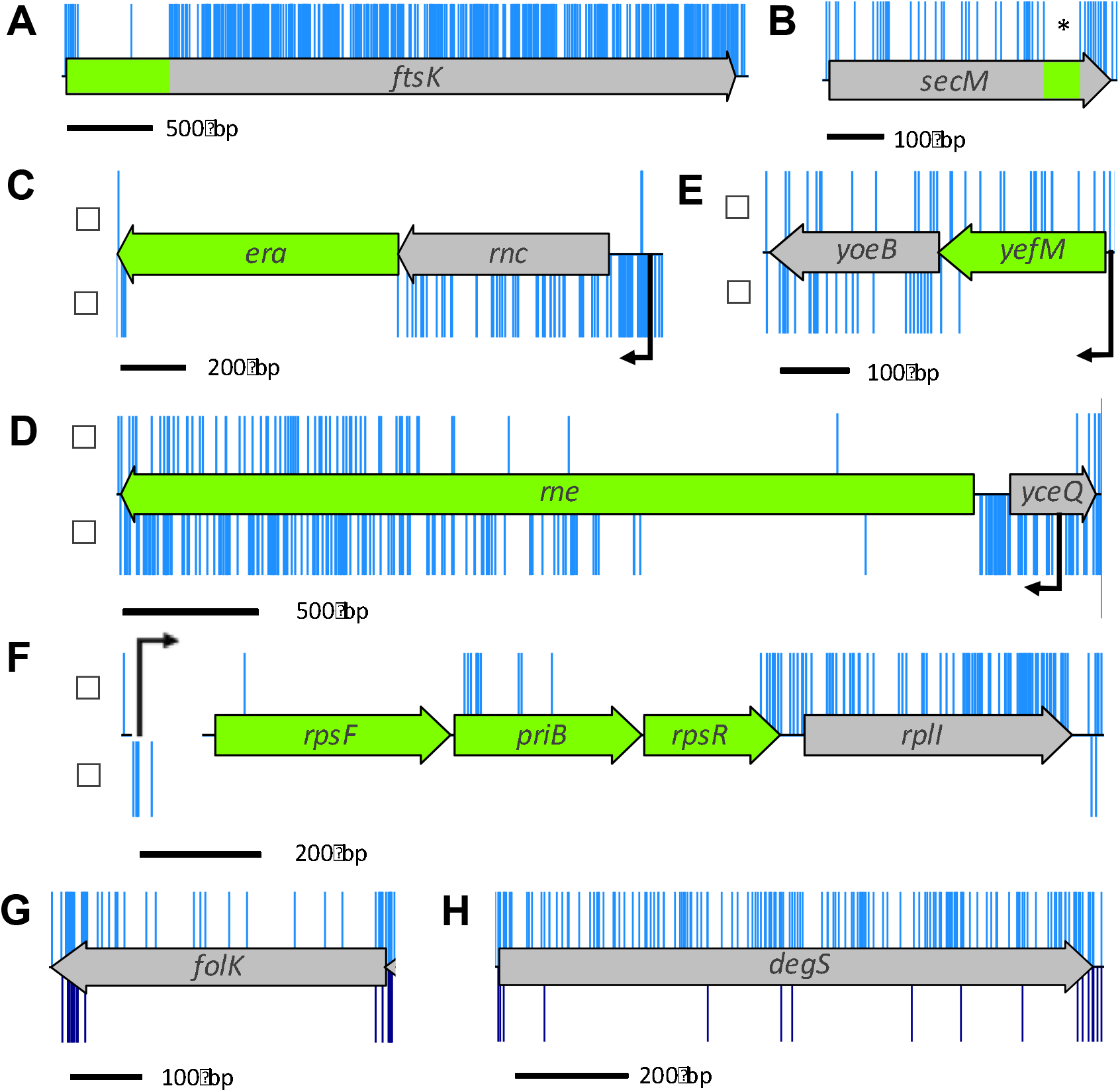
Insertion profiles of discrepent genes between datasets. (A) The *ftsK* gene codes for an essential protein in which only part of the protein is required for its essential function. Such genes have a high insertion index score and consequently would not have been identified by automated statistical analysis. (B) *secM* contains a window (*) of 66 bp in which there were no transposon insertions. This feature is discussed in the text. (C-F) Genes with transposon insertions in only one orientation. The α- and β-orientation of the transposon is depicted above and below the CDS track respectively, native promoters are shown in black. (G and H) Many transposon insertions were found along the full length of *folK* and *degS* (shown in blue in the upper track of the figure). However, most of these insertion mutants were lost during outgrowth (lower track below, dark blue).

The second group contains genes with transposon insertion sites throughout the CDS but which have an IFR that passes the significance threshold for essentiality. For example, there is a small IFR within the coding sequence of *secM* of 66 bp (Fig. 3B, Table S3). The *secM* gene is located upstream of the essential gene *secA*. These genes are co-transcribed and also co-translated, and *secM* is known to contain a translational stop sequence that interacts with the ribosomal exit tunnel to halt translation, acting as a translational regulator for *secA*. Specific mutations within the translational stop sequence are lethal unless *secA* is complemented by expression from a plasmid (26). The dependence of *secA* translation on the *secM* CDS would explain the Keio classification as ‘essential’. However, the IFR within *secM* does not fully correspond with the translation stop sequence, suggesting that there is more to be learnt about the translational linkage between the two proteins.

Other researchers have used different approaches to minimize false classification of essential genes during statistical analysis of the insertion profiles by applying a sliding window, quantifying the mean distance between insertions per gene, or variations of truncating the CDS, such as excluding the 3′ end, analyzing only the first 60% of the CDS or analyzing the central 60% of the CDS (18, 19, 27–31). However, window analysis may overlook genes such as *secM* and analyzing only the first 60% of the CDS would overlook genes such as *ftsK*.

We suggest that the algorithmic approach used here is a more appropriate method for identifying essential chromosomal regions in a sufficiently dense library. However, we see a number of IFRs >45 bp throughout the genome within non-essential genes, suggesting that our null model of random insertions is not capturing the full structural detail of transposon insertion propensity. This suggests our modelling approach is not based on a perfect representation of biological reality and needs further refinement.

#### Polar insertions

A common feature when creating insertion mutants is the introduction of off-target polar effects where expression of adjacent genes are disrupted by the insertion. To mitigate against such polar effects we designed a cassette that enabled both transcriptional and translational read-through in one direction only. To confirm that transcriptional and translational read-through emanates from the transposon, the transposon was cloned in both orientations and in all three reading frames upstream of the *lacZ* gene in transcription and translation expression vectors pRW224 and pRW225, derivatives of pRW50 (32, 33). Transcriptional read-through was confirmed for one orientation of the transposon, consistent with transcriptional read-through from the chloramphenicol resistance cassette into the downstream disrupted CDS (Fig. 4). Translational read-through was identified for 2 of the 3 open reading frames that coincided with AUG and GUG start codons in the inverted repeat at the end of the transposon. More β-galactosidase activity was obtained from the construct in which the AUG codon was in frame than when the GUG codon that was in frame, confirming that translation was initiated more strongly from the AUG codon. Therefore, transcription is initiated from within the transposon and translation is initiated from within the inverted repeat. This allows transcription and translation of downstream essential regions, even from within a CDS. Such events can be identified by determining to which DNA strand the sequencing data maps (Fig. 4C).

**Figure 4.**
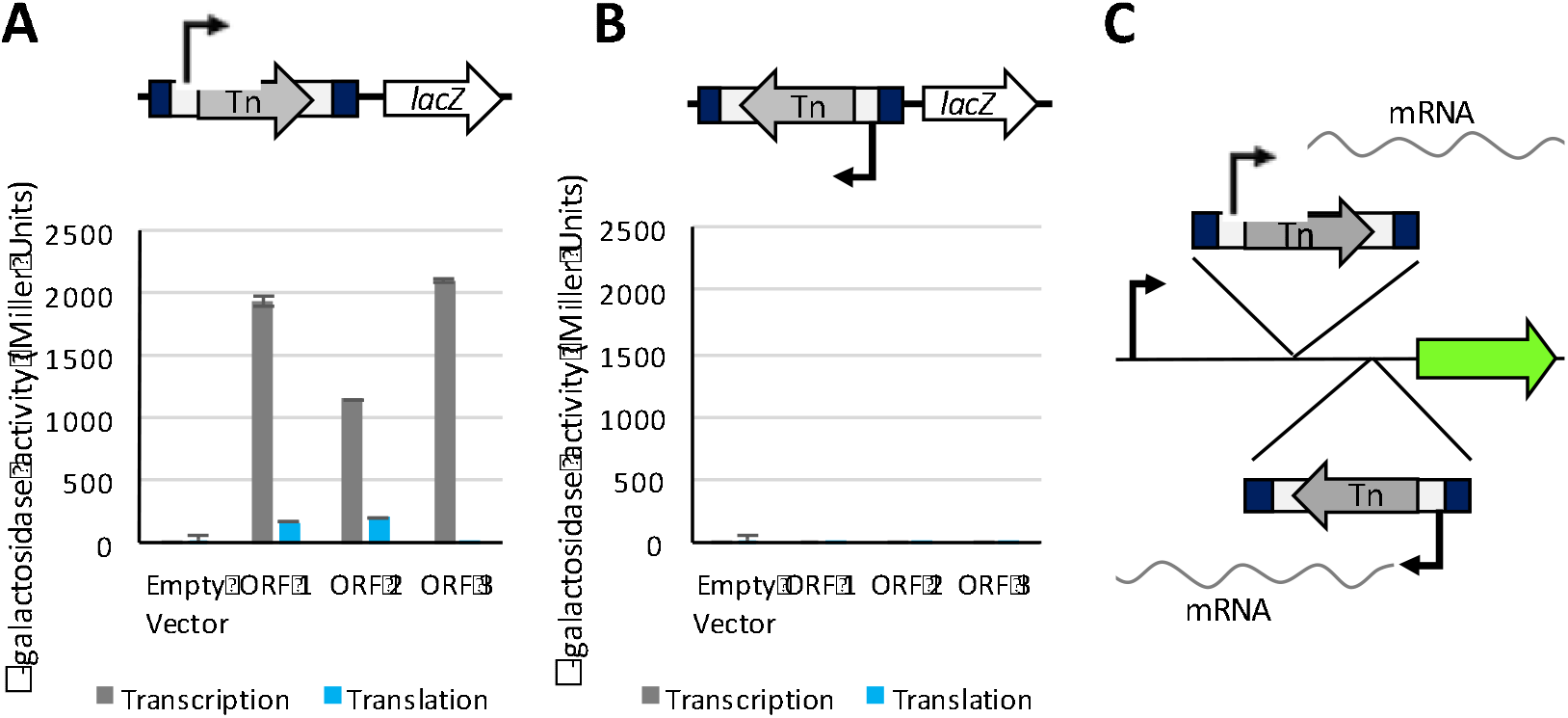
Transcription and Translation initiation from within the transposon. The full length mini-Tn*5* transposon was cloned into expression vectors pRW224 and pRW225 upstream of the *lacZ* gene, in each orientation, for all three open reading frames (ORF). Vector pRW224 retains a RBS for *lacZ* but no promoter, vector pRW225 has no promotor or RBS upstream of *lacZ*. Vectors pRW224 and pRW225 can be used to detect transcriptional and translational activity respectively. β-galactosidase activity was measured in triplicate for 3 technical replicates, the mean value was plotted with error bars showing the standard deviation between replicates. (A) Transcriptional read-through was confirmed for one orientation of the transposon, consistent with the orientation of the chloramphenicol gene. Translational read-through from the mini-Tn*5* transposon was confirmed for 2 out of 3 open reading frames, consistent with GUG (ORF1) and AUG (ORF2) start codons in the transposon inverted repeat. (B) No transcriptional or translational read-through was detected for the opposite orientation of the transposon. (C) Schematic representing the orientation of transposon insertions. The α-orientation of the transposon (upper track) corresponds with the chloramphenicol cassette oriented left to right. The β-orientation (lower track) corresponds with transposon insertions in the opposite direction.

Analysis of our data reveals a number of chromosomal regions with insertions in only one orientation. Such insertion profiles can offer insight into transcriptional regulation of genes when considered in conjunction with neighbouring genes. For example, the gene *rnc* is located in an operon upstream of the essential gene, *era*. Only mutants with transposons that maintain downstream transcription of *era* are viable (Fig. 3C). Baba *et al*. categorized *rnc* as essential (34). However, in the case of the Keio library, construction of an *rnc* deletion mutant would disrupt the ability of the native promoter to drive downstream expression of the essential *era* gene, resulting in apparent lethality. Similarly, in both the Keio and PEC databases *yceQ* is listed as essential but we observed many insertions in *yceQ*, but in only one orientation (Fig. 3D). The gene is located upstream to the essential gene *rne* and is divergently transcribed. The promoter for *rne* is positioned within *yceQ* (35, 36) and deletion of *yceQ* would remove the promoter for *rne* resulting in an apparent lethal effect. Our data reveal that while *era* and *rne* are essential, *rnc* and *yceQ* are not essential.

Like *rnc* and *yceQ*, several of the anti-toxin genes are reported to be essential in the Keio library but not in our dataset or the PEC database (Table S3). Antitoxins are required only if the corresponding toxin gene is functional. One example is *yefM*. We observed a substantial number of insertions in one orientation. Unlike *rnc* and *yceQ* where insertions maintained downstream expression, in the case of *yefM* the opposite is true; insertions that disrupt expression of the antitoxin but maintain downstream expression of the downstream toxin (*yoeB*) are lethal (Fig. 3E). Scrutiny of our data in this manner reveals these genes are essential.

Another example of insertion bias is observed in a number of genes at the 3′ end of a transcript, such as *rplI* (Fig. 3F). While *rplI* is not reported as essential, it is worth noting because insertions restricted exclusively to one orientation within the gene cannot be explained by the positional context between an essential gene and promoter. One possible explanation for this observation is that transcription promoted from the transposon produces an antisense RNA that inhibits expression of an essential gene. Insertion bias, irrespective of the underlying cause, can result in false classification of genes when quantifying insertion index scores as these genes have half as many insertions relative to the rest of the genome. As such, these insertion profiles are to be considered when analyzing data with autonomous statistical approaches.

#### Conditionally essential genes

In addition to the scenarios listed above, certain genes present challenges for binary classification of essentiality. For example, the genes might code for a protein that is essential at a specific phase of growth or for growth under certain environmental parameters such as temperature or nutrient availability. Our data reveal a range of these conditionally essential genes. For instance, the Keio and PEC databases list *folK* as essential whereas we detected multiple insertions within *folK* (Fig. 3G). Loss of *folK* disrupts the ability of the bacterium to produce folate, which is an essential metabolite. However, supplementation of the medium with folate abrogates the requirement for folate biosynthesis. In addition to *folK*, the Keio and PEC databases report *degS* as essential. In our dataset *degS* has a high density of insertions throughout the CDS suggesting that *degS* is not essential for growth on an agar plate (Fig. 3H). Consistent with this, there is substantial literature showing that *degS* mutants can be isolated, but they either lyse in the stationary phase of growth or rapidly accumulate suppressor mutations (37–40).

The conditional essentiality of such mutants can be tested by growing the transposon library in liquid broth. One would expect mutants lacking *degS* will lyse and that *folK* mutants will be outcompeted as the limited folate available in the medium is depleted. To test these scenarios, two independent samples of the transposon library were grown in LB at 37 °C for 5 to 6 generations to an OD_600_ of 1.0 and were then sequenced. These samples, LB1 and LB2, resulted in 5,908,163 and 6,403,324 sequences of which 5,201,711 (88.04%) and 5,382,477 (84.06%), respectively, were mapped to the BW25113 genome (Table 1). Insertion index scores were calculated as before (Table S4). As there was a high correlation coefficient of 0. 97 between the gene insertion index scores of each technical replicate (Fig. 1C), the data were combined to give a pool of 10,584,188 sequences. Scrutiny of our data revealed substantially fewer *degS* and *folK* mutants after growth in LB, supporting our hypothesis that they are conditionally essential (Fig. 3G and H). Other genes showing similar fitness costs can be identified in the LB outgrowth dataset (Table S4).

#### Errors in library construction

The difficulty in classifying a gene as essential through deletion analysis is the dependence on a negative result to inform classification. Thus, failure to knock out the gene may result in the false classification of a gene as essential. For example, the Keio database originally reported *mlaB* (*yrbB*) as essential. However, our data demonstrate that *mlaB* is non-essential and this is supported by the literature (41, 42). We have observed similar outcomes for several other genes (Table S3). The reason why knockouts of these genes were not obtained in the construction of the Keio library is unknown.

In addition to the false positive outcomes described above, we noted several instances of false negative results within the Keio library database. For example, both our TraDIS data and the PEC database identified 18 genes as essential that are reported as non-essential in the original Keio database (Table S2). Subsequently, Yamamoto *et al*. (34) demonstrated that for 13 of these mutants the target gene was duplicated during construction of the Keio library resulting in a functional protein; these genes are almost certainly essential. Another difficulty that arises when targeting essential genes for mutagenesis is the potential to select for mutants with a compensatory mutation elsewhere in the genome. Our data revealed *hda* is an essential gene but it is classified as non-essential in the Keio database. Since the initial description of the Keio library, *hda* has been reported to be essential but *hda* mutants rapidly accumulate suppressor mutations that restore viability (43–45). We hypothesise that this is an explanation for the observed essentiality of some genes in the TraDIS dataset that were described as non-essential by others (Table S3). These effects may arise when creating TraDIS libraries but the effects are masked by the large number of mutants in the population.

Similarly, in the PEC library, where insertion density is low, essential genes with an insertion in a non-essential region of the gene will be falsely classified as non-essential when relying on single insertion mutants to inform essentiality. An example of this false negative classification in the PEC database is *tadA* (Table S3). The TadA protein is a tRNA-specific deaminase and its essentiality is reported in the Keio database and our dataset, and is supported by the literature (46). The PEC database reports a single insertion site within the extreme 3′ end of the *tadA* gene.

We have identified a range of underlying causes behind dataset discrepancies and highlight that there are numerous possible insertion profiles for an ‘essential’ gene. As such, it is important to note that no single statistical method, to our knowledge, would fully identify every essential gene and that manual inspection of data is crucial.

### Genes identified as essential only by TraDIS

There are 81 genes identified as essential using our insertion index data, which are not reported as essential in the Keio or PEC databases (Table 2). These genes fall into two groups, those with no insertions and the remainder with insertions in the CDS. The first group are most likely to be essential. For example, *rpsU* is essential in our data and has been described as essential by others (Fig. 5A; 47). However, in the Keio library there is a duplication event, which gives rise to a mutant that produces a functional protein (34).

**Figure 5.**
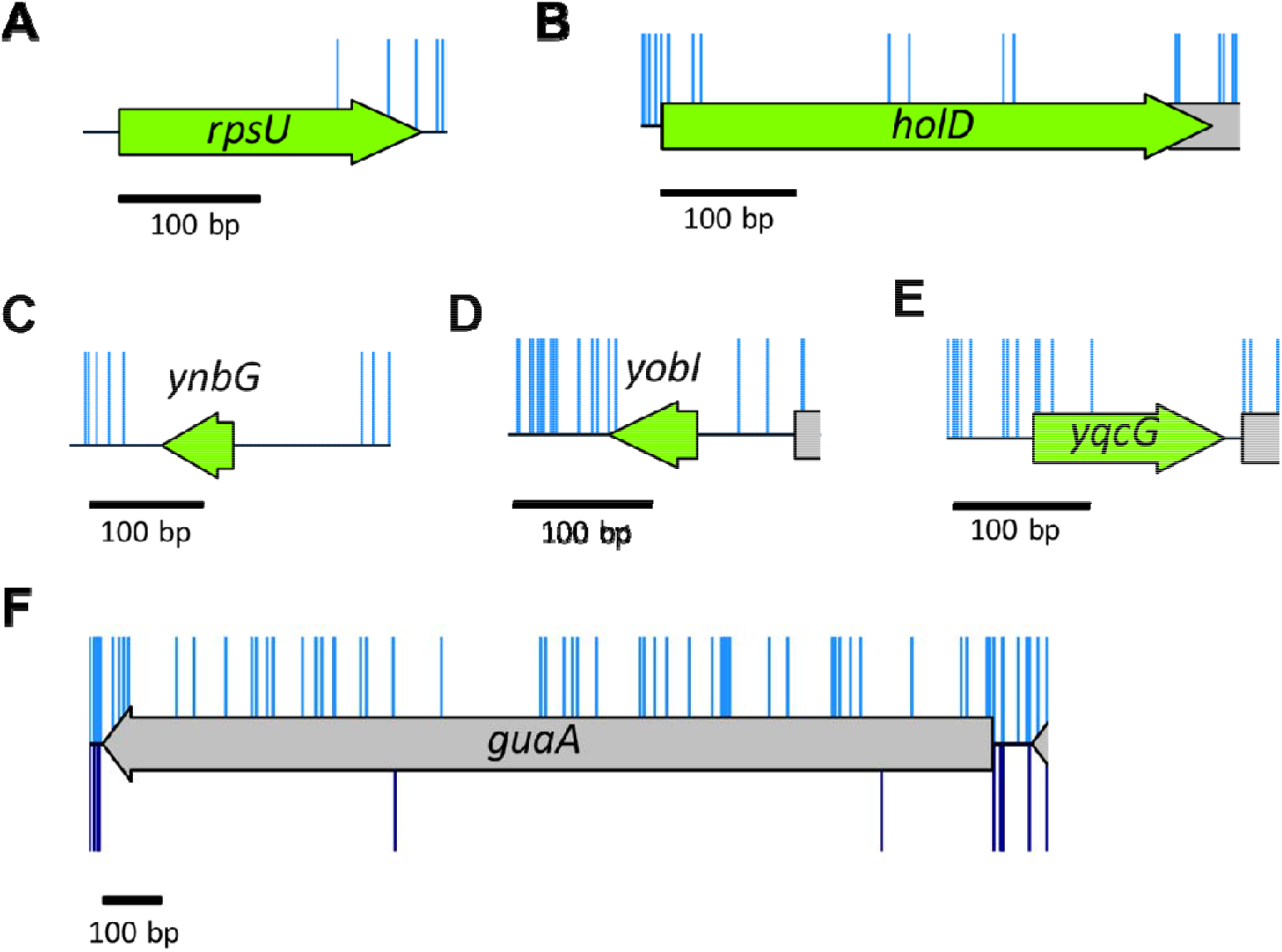
Essential genes unique to our data. (A-E) There are very few or no insertions within these genes in our input library (blue). (A & B) Low insertion frequency and literature support classification of these genes as essential. (C, D & E) Recently annotated genes with few or no insertions. Our data suggest these are potentially essential or important for growth. (F) The gene *guaA* has a sufficiently low insertion index score to be classified as essential after initial statistical analysis (blue, above). Following outgrowth, there are few *guaA* mutants (dark blue, below), consistent with literature reports that *guaA* mutants have a growth defect.

**Table 2.**
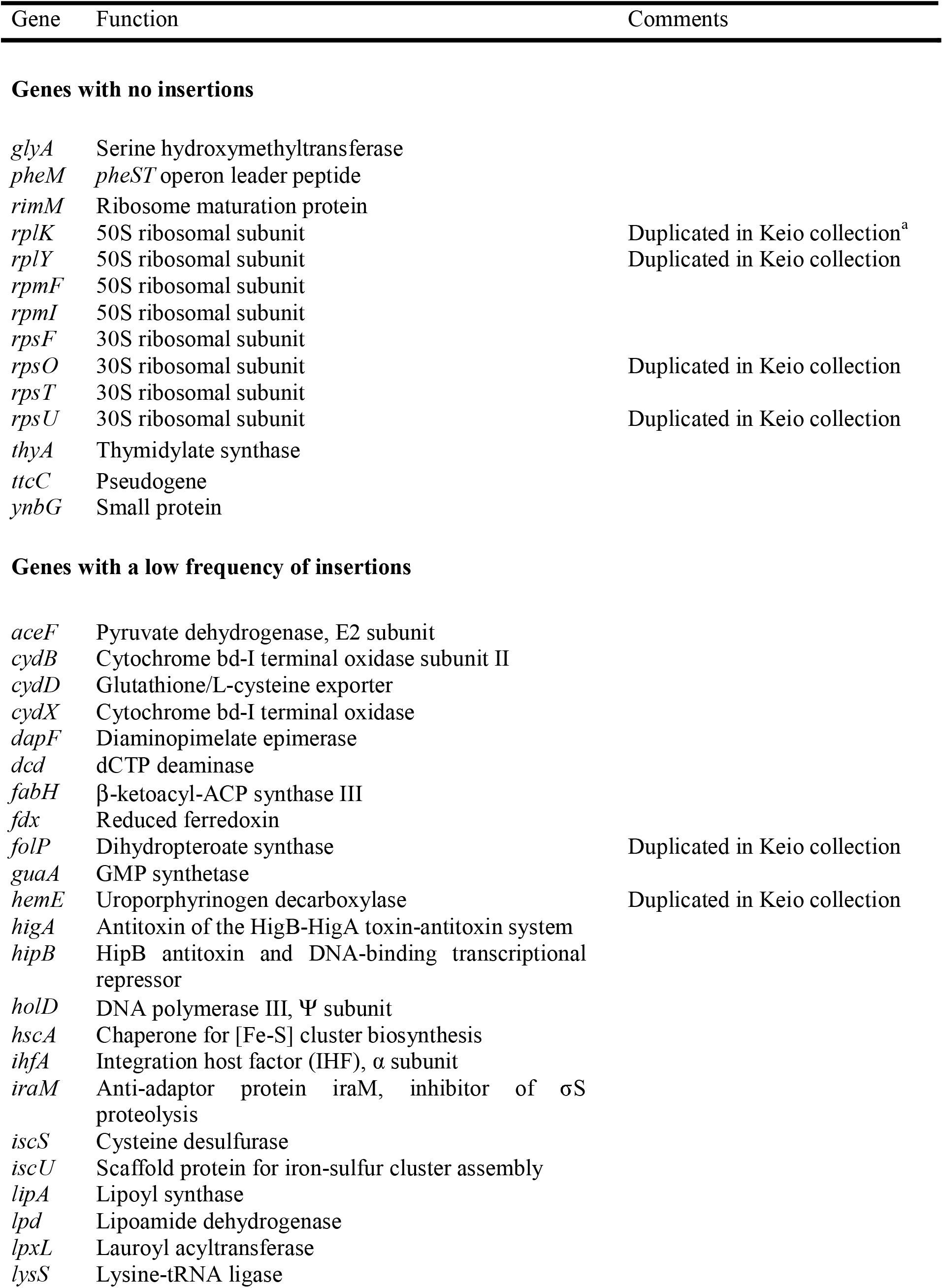
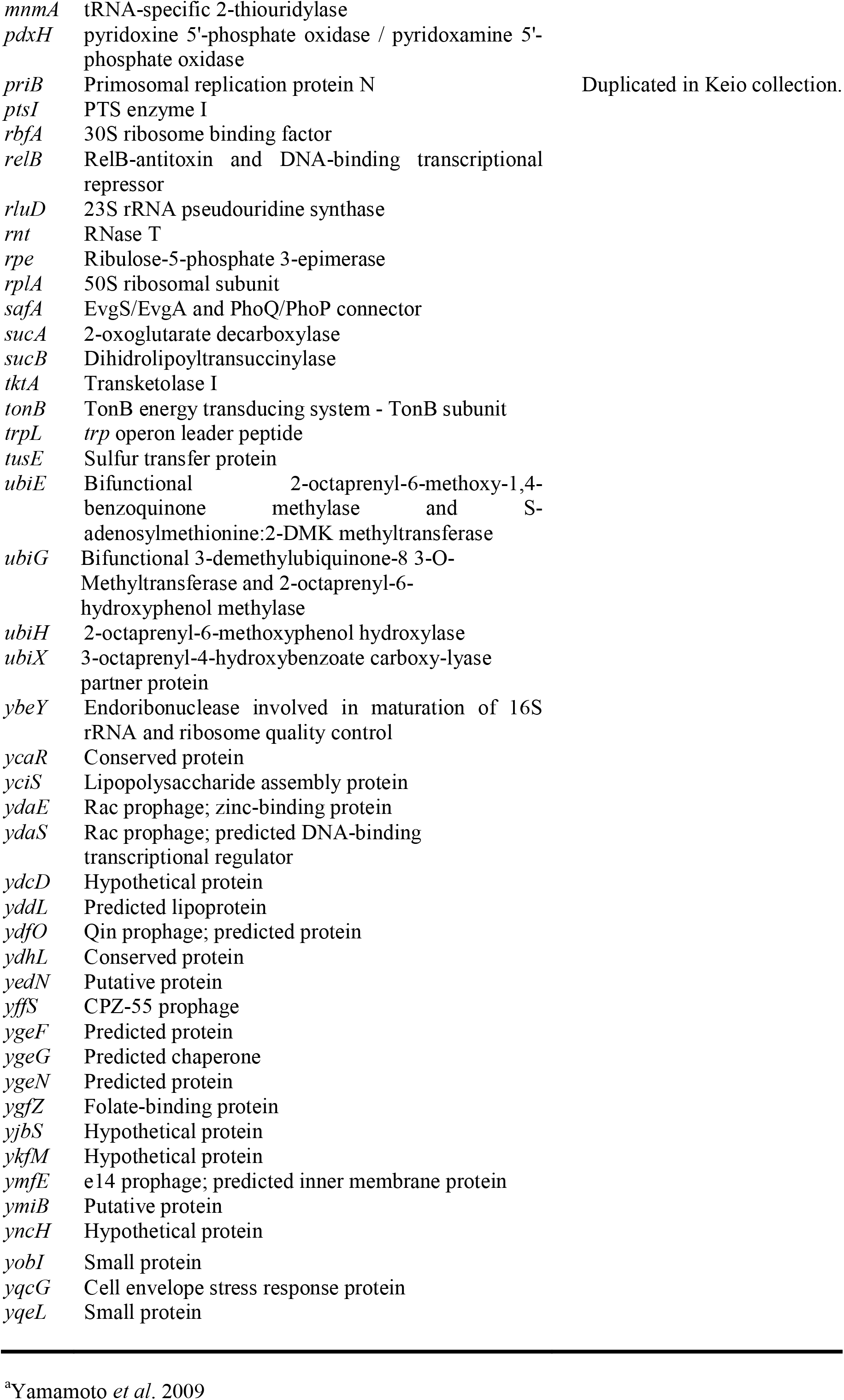
Genes with low insertion index scores identified by TraDIS only

Scrutiny of our data for the remaining genes reveals there are additional essential genes with a low frequency of insertions. For instance, *holD* has been described in the literature as an essential gene (48). Our data support that finding (Fig. 5B, Table S1). However, *holD* mutants are available in the Keio collection. The demonstration by Durand *et al*. and others (48–50) that *holD* mutants accumulate extragenic suppressor mutations at high frequency may explain why these mutants are considered non-essential in the Keio database and why we observe a low frequency of insertions in our experiments.

A number of the genes unique to our analysis were not identified as essential in the Keio collection or PEC database simply because they are not included in either of these datasets. This is in part because the Keio collection of knockout mutants was based on available annotation data at the time (51). For example, the identification and location of *ynbG, yobI* and *yqcG* was published only in 2008 (52). These genes show very sparse or no transposon disruption in our data and consequently, these genes are potentially essential (Fig. 5C, D and E). Further validation studies would be required to confirm this.

As mentioned previously, over-reporting of essential genes may occur when non-essential genes have low insertion index scores. Such low insertion index scores may arise due to attenuated growth. An example of gene mis-classification because mutation results in a fitness cost and attenuated growth is *guaA*. The low insertion index score results in it being classed as essential despite having many insertions. The fitness effect was confirmed by growing the library in LB, as such mutants are outcompeted (Fig. 5F), and the literature supports the fact that this gene is not essential and has an altered growth rate (53).

### High resolution features within a TraDIS dataset

Manual inspection of a TraDIS dataset can reveal additional information that might go unnoticed in a high throughput analysis pipeline. A common observation from this and previous detailed analysis of data from saturated transposon libraries is the ability to determine, at the bp level of resolution, the boundaries of essential regions within a gene. An example of an essential gene with a dispensable 3′ end is *yejM* (*pbgA* in *Salmonella* Typhimurium). Only the 5′ end of the CDS is essential, up to and including codon 189, which corresponds with 5 transmembrane helices of the protein structure; the C terminus of the protein is a periplasmic domain that is dispensable for viability (Fig. 6A; 54–56). Our TraDIS data revealed insertions in codons 186 and 189. Analysis of the transposon orientation at these points revealed that they corresponded with the same transposon insertion location but, due to the 9-bp duplication introduced by the transposon, in different transposon orientations. The introduced transposon sequence maintains codon 189, completely consistent with previously reported results (54, 56).

**Figure 6.**
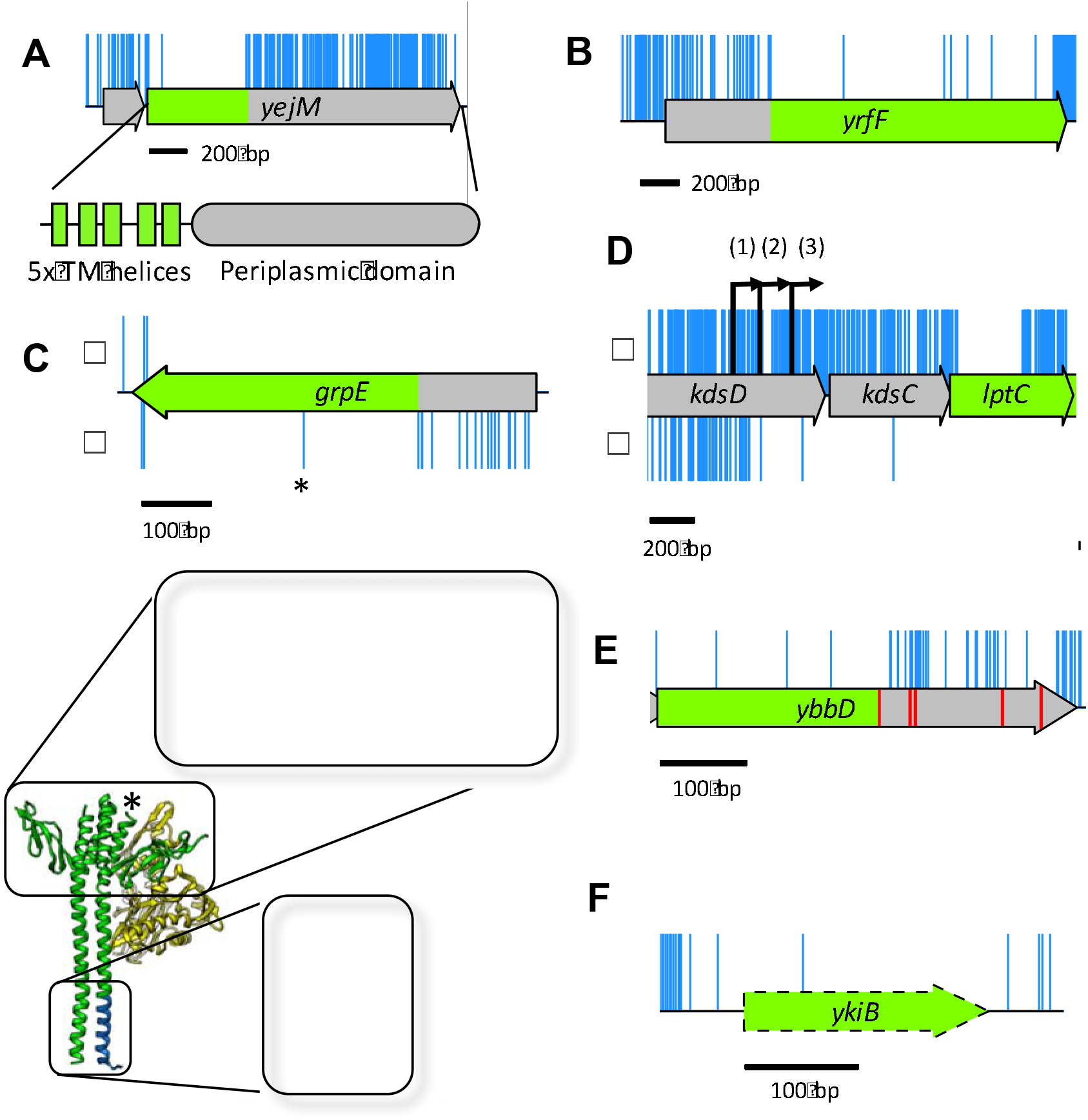
Additional features identified through detailed analysis of high-resolution insertion data. (A) Insertions within the CDS but not along the full length correspond with a non-essential periplasmic domain. The 5′ end of the CDS has no insertions and corresponds with 5 essential transmembrane domains of YejM. (B) Insertions within *yrfF* suggest a dispensable 5′ domain. (C) The *grpE* gene tolerates transposon insertions in the 5′ end of the CDS (blue), but only in the orientation that maintains expression of the downstream protein (lower track, β-orientation). The GrpE protein forms a dimer (green) which interacts with DnaK (yellow). Transposon insertions in specific regions of the protein do not disrupt GrpE interaction with DnaK (blue). An additional, single, insertion point in the center of the *grpE* CDS (*) maps back to a turn between 2 helices of the GrpE protein. The data reveal dispensable sections of the GrpE protein and boundaries in secondary structure. (D) Insertions immediately upstream of *lptC* have an insertion orientation bias. Only insertions that maintain expression of *lptC* are tolerated within *kdsC* (α-orientation). The gene *lptC* has 3 promoters (‘1’, ‘2’ and ‘3′), the insertion boundary indicate that promter ‘2’ is the essential promoter. (E) Pseudogene *ybbD* contains many more insertions after the first stop codon (red), suggestive that the truncated CDS may still be functional and essential. (F) The pseudogene *ykiB* is not annotated in the BW25113 genome (CP009273.1) and has a single insertion within the CDS.

In addition, as a result of our transposon design, a further feature of our TraDIS data is the identification of genes with dispensable 5′ ends. An example of this is *yrfF* which encodes an inhibitor of the Rcs stress response (Fig. 6B; 57, 58). This phenomenon, while less well covered in the literature, is not surprising given that Zhang *et al*. report equal likelihood of a required intragenic region residing at the 5′ or 3′ end of a gene, albeit in *Mycobacterium tuberculosis* (31). These mutants will only be viable if the remaining CDS can be translated into a functional product and one would expect to find an orientation bias where the transposon drives downstream transcription and translation of the essential region.

Interestingly, inspection of our data revealed essential genes with isolated insertions within the coding sequence. An example of this is *grpE*. The *grpE* gene codes for the essential nucleotide exchange factor that forms a dimer and interacts with DnaK/J complex (59). The insertion occurs only in the orientation that maintains expression of the remaining CDS (Fig. 6C). Mapping of the site of transposon insertion onto the previously determined protein structure of GrpE indicated the insertion occurred within part of the gene encoding a flexible linker between two α-helices (Fig. 6C). This suggests that a functional GrpE can be expressed as two separate essential domains, and that our TraDIS library has an unprecedented sub-CDS level of resolution to demarcate changes in the protein secondary structure.

Another fine-resolution mapping feature of our transposon data is the identification of the promoter position for essential genes, as also previously reported by Christen *et al*. (8). An example of this is the promoter for *lptC*, located within the *kdsD* gene. Three promoters (kdsCp3, kdsCp2 and kdsCp1) have been identified within the *kdsD* gene (60, 61). However, our data show that insertions that maintain expression of *lptC* are tolerated within the *kdsD* gene up to kdsCp2; insertions stop short just before the kdsCp2 −35 consensus sequence, excluding a single insertion between the −35 and −10, and a single insertion further downstream (Fig. 6D). These results suggest that kdsCp3 is dispensable and at least kdsCp2 is required for adequate expression of *lptC*. As in the case of *grpE* above, this is another example of the unprecedented level of genetic detail that can be obtained from this high throughput method.

Finally, within our data we observed a number of transposon-free sections that do not correspond with the annotated features of our genome. This can occur when a start codon is mis-annotated (8), or translation might initiate at a secondary start codon downstream of the transposon insertions. Alternatively, a pseudogene annotation may extend beyond the first stop codon. One example of this is *ybbD*, a pseudogene classified as non-essential in our dataset. However, the annotation of *ybbD* in the BW25113 genome extends beyond the first stop codon, whereas in other genomes it does not, and the region from the methionine translation start codon to the first stop codon passes the threshold for essentiality (Fig. 6E). In addition, we observed a transposon free region corresponding to *ykiB* (Fig. 6F). This gene is annotated in W3110 but not in BW25113, despite the fact the nucleotide sequence is identical. Our data suggest that these genes have a significant role in viability or growth but further investigation is required to test this. These examples highlight the importance of having a fully annotated and curated reference genome for mapping data. However, even the highly-studied K-12 genome with its wealth of annotation information retains some as yet unexplained transposon free regions. Thus, TraDIS can help identify regions of genomes where annotation is incorrect or incomplete.

### Conclusion

In summary, the current study has identified 248 genes that are essential for *E. coli* K-12 to survive in standard laboratory conditions (Table S2). We have shown why different conclusions have been drawn from transposon mutagenesis data and gene deletion studies. Essential genes that contain both essential and non-essential regions will statistically appear non-essential if judged only on insertion index scores. We have demonstrated the importance of visual analysis to avoid automation bias in designating genes as essential or non-essential. We have also identified genes incorrectly designated as essential because of failure to recognize polarity effects on a downstream essential gene in the same transcription unit. We also report potential new essential genes and discuss the use of transposon sequencing for fine-resolution mapping of features across the genome. Importantly, TraDIS data are a valuable resource that can be re-inspected following the discovery of new features within a given genome. Finally, our data reveal that there is more to be understood about genome structure and organization, which further coupling of modelling and experimental approaches will help to elucidate.

## MATERIALS AND METHODS

### Strains and plasmids

*E. coli* K-12 strain BW25113, the parent strain of the Keio library, was used for construction of a transposon library. The strain has the following genotype: *rrnB3* Δ*lacZ4787 hsdR514* Δ(*araBAD*)*567* Δ(*rhaBAD*)*568 rph-1* (62). The transposon mutant library was constructed by collaborators from Discuva, Cambridge, following a method described for *Salmonella* Typhi (4). The main differences were that a mini-Tn*5* transposon coding for chloramphenicol resistance cassette resistance was used. This was amplified by PCR from the *cat* gene of the plasmid vector pACYC184 (63) using oligonucleotide primers incorporating the Tn*5* transposon mosaic ends. Transposomes were prepared using Tn*5* transposase (Epicentre, Madison, WI, USA), and these were introduced into *E. coli* K-12 strain BW25113 by electrotransformation. Transposon mutants were selected by growth on LB agar supplemented with chloramphenicol. Approximately 5.6 million colonies representing an estimated 3.7 million mutants were pooled and stored in 15% glycerol at −80 °C.

### Media and growth conditions

DNA was extracted from two samples of the transposon library glycerol stock to generate TraDIS data referred to as NTL1 and NTL2 in the text. In addition, DNA was extracted from two independent cultures, LB1 and LB2, of the library grown in Luria broth (LB: 10 g tryptone, 5 g yeast extract, 10 g NaCl) and grown for 5 to 6 generations at 37 °C with shaking until OD_600_ = 1.0.

### Beta-galactosidase assay

β-galactosidase assays were used to measure the activity of transposon::*lacZ* fusions. The transposon was cloned in each orientation, for all three open reading frames, into transcription and translation-assay vectors pRW224 and pRW225 (33). Strains carrying the transposon::*lacZ* fusions were grown overnight at 37 °C with aeration in LB supplemented with 35 μg/ml tetracycline (Sigma). The density of the overnight culture was determined by measuring OD_650_ and then used to sub-culture into 5 ml LB and incubated at 37 °C with aeration until the mid-exponential phase of growth (OD_650_ of 0.3-0.5). Each culture was lysed by adding 100 μl each of toluene and 1% sodium deoxycholate, mixed by vortex for 15 s and aerating for 20 min at 37 °C. The β-galactosidase activity of each culture was assayed by addition of 100 μl of each culture lysate for three technical replicates to 2.5 ml Z buffer (10 mM KCl, 1 mM MgSO_4_.7H_2_O, 60 mM Na_2_HPO_4_, 30 mM NaH_2_PO_4_.2H_2_O supplemented with 2.7 ml β-mercaptoethanol per litre of distilled water, adjusted to pH 7) supplemented with 13 mM 2-nitrophenyl β-D-galactopyranoside (ONPG; Sigma). The reaction was incubated at 37 °C until a yellow colour had developed, after which the reaction was stopped by adding 1 ml 1 M sodium carbonate. The absorbance of the reaction at OD_420_ was measured and β-galactosidase activity was calculated in Miller units.

### TraDIS Sequencing

Harvested cells were prepared for sequencing following an amended TraDIS protocol (4, 8, 9). Genomic DNA was isolated using a Qiagen QIAamp DNA Blood Mini Kit, according to the manufacturer’s specifications. DNA was quantified and mechanically sheared by ultrasonication. Sheared DNA fragments were processed for sequencing using NEB Next Ultra I kit. Following adaptor ligation, a PCR step was introduced to enrich for transposon-containing fragments, using a forward primer specific for the transposon 3′ end and a reverse primer specific for the adapter. After PCR-purification, an additional PCR prepared DNA for sequencing through the addition of Illumina-specific flow cell adaptor sequences and custom inline index barcodes of variable length in the forward primers. The purpose of this was to increase indexing capacity while staggering introduction of the transposon sequence to increase base diversity during sequencing. Samples were sequenced using Illumina MiSeq 150 cycle v3 cartridges, aiming for an optimal cluster density of 800 clusters per mm^2^.

### Sequencing analysis

Raw data was collected and analyzed using a series of custom scripts. The Fastx barcode splitter and trimmer tools, of the Fastx toolkit, were used to assess and trim the sequences (64). Sequence reads were first filtered by their inline indexes, allowing no mismatches. Transposon similarity matching was done by identifying the first 35 bp of the transposon 5′ end in two parts: 25 bases (5′ to 3′) were matched allowing for 3 mismatches, trimmed, and then the remaining 10 bases matched allowing for 1 mismatch, and trimmed. Sequences less than 20 bases long were removed using Trimmomatic (65). Trimmed, filtered sequences were then aligned to the reference genome *E. coli* BW25113 (CP009273.1), obtained from the NCBI genome repository (66). Where gene names differed between databases, the BW25113 annotation was used. The aligner bwa was used, with the mem algorithm (0.7.8-r455; 55). Aligned reads were filtered to remove any soft clipped reads. The subsequent steps of conversion from sam files to bam files, and the requisite sorting and indexing, were done using samtools (0.1.19-44428cd; 68). The bedtools suite was used to create bed files which were intersected against the coding sequence boundaries defined in general feature format (.gff) files obtained from NCBI (69). Custom python scripts were used to quantify insertion sites within the annotated CDS boundaries. Data were inspected manually using the Artemis genome browser (70).

### Essential gene prediction

The frequency of insertion indices was plotted in a histogram using the Freedman–Diaconis rule for choice of bin-widths (Figure S1). Using the R MASS library (<u>http://www.r-proiect.org</u>), an exponential distribution (red line) was fitted to the left, “essential” mode (i.e. any data to the left of the trough in Figure S1); a gamma distribution (blue line) was fitted to the right, “non-essential” mode (i.e. any data to the right of the trough). The probability of a gene belonging to each mode was calculated and the ratio of these values was used to calculate a log-likelihood score. Using a 12-fold likelihood threshold, based on the log-likelihood scores, genes were assigned as “essential” if they were 12 times more likely to be in the left mode than the right mode, and “non-essential” if they were 12-times more likely to be in the right mode (9). Genes with log-likelihood scores between the upper and lower log_2_(12) threshold values of 3.6 and −3.6 respectively were deemed “unclear”. A threshold cut-off of log_2_(12) was chosen as it is more stringent than log_2_(4) (used by Langridge *et al*.; 4), and consistent with analysis used by Phan *et al*. (9).

### Essential gene lists

The Keio essential gene list is composed of the original essential genes minus three ORFs, JW5190, JW5193 and JW5379, as they are not annotated within MG1655 and are thought to be spurious, giving a final list of 300 genes (1, 71). The PEC dataset is composed of the 300 genes listed as essential for W3110 (2). The lists of essential genes were compared using BioVenn (72).

### Statistical Analysis

For details of the statistical analysis, see Supplementary materials and methods and Figures S1 and S2.

## ACKNOWLEDGEMENTS

We thank Discuva Ltd. for providing some of their large transposon mutant library. We thank N. Loman and J. Quick for help with optimization of our MiSeq protocol. We thank the authors of Langridge *et al*. (2009) for kindly supplying their R code for essential gene prediction. We thank Tony Hitchcock and Steve Williams for their support, and Cobrabio for their contribution to the University of Birmingham Elite PhD studentship. Lastly, we thank M.J. Collingwood and R.W. Meek for their generous help with drawing figures.

## Supplementary Figures

**Figure S1.**
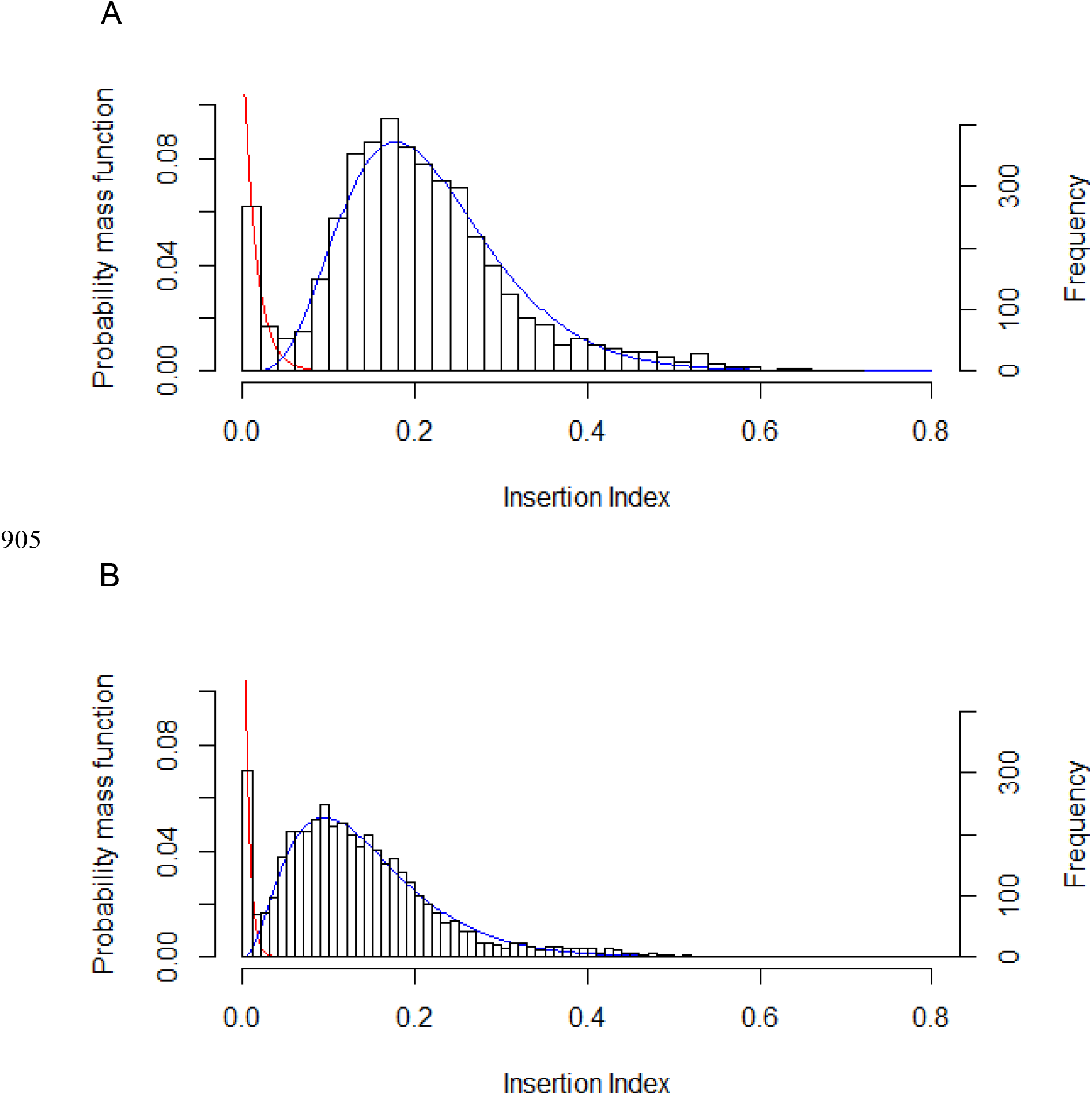
Frequency distribution of insertion index scores. The insertion index score for each coding sequence was calculated as the number of insertions per CDS divided by the CDS length in bp to normalize for gene length. The frequency of insertion index scores was plotted for both NTL (A) and LB data (B) and both followed a bimodal distribution. An exponential distribution model was fitted to the left mode that includes essential genes (red) and a gamma distribution model was fitted to the right, non-essential mode (blue). For a given insertion index score, the probability of belonging to each mode was calculated and the ratio of these values termed the log-likelihood score. A gene was classified as essential if its log-likelihood score was less than log_2_(12) and was therefore 12 times more likely to belong to the red mode than the blue mode

**Figure S2.**
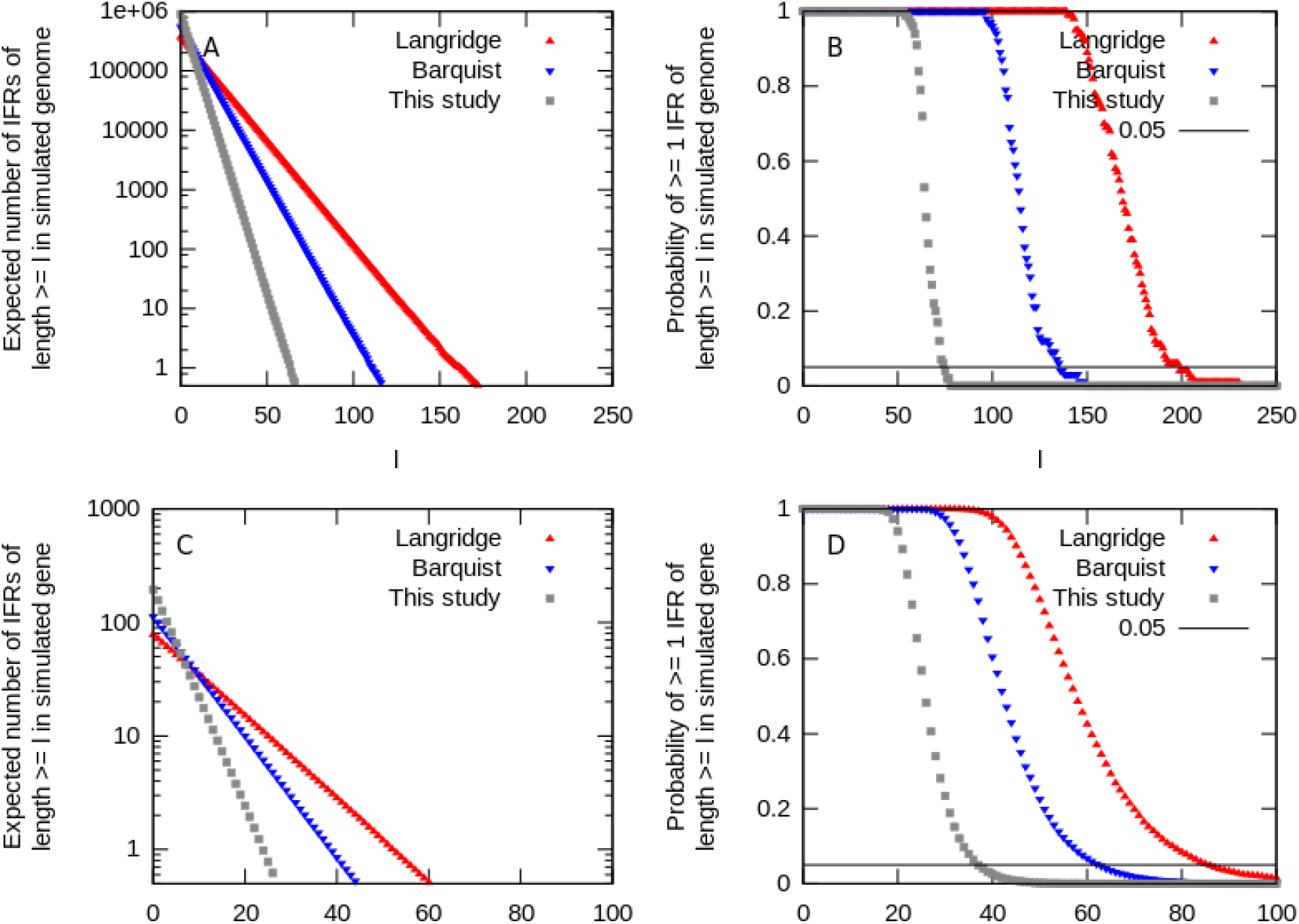
Simulation of random insertion events for rigorous statistics under the null model of random insertion. Simulation of random insertion events for rigorous statistics under the null model of random insertion. (A) The expected number of insertion-free regions (IFRs) of length l or above found in a genome of length G under the null model of N random non-coincident insertions. G and N are taken to correspond to the statistics in the studies plotted. This expectation is taken over 100 instances of the null model simulation. (B) The (related) probability of at least one IFR of length l or above occurring anywhere in the simulated genome. (C, D) The same calculation over a simulated gene of length g = 1000, using 10^5 simulation instances.

## Supplementary tables

**Table S1. Essential genes identified by TraDIS as essential**

**Table S2. Comparison of essential genes between datasets**

**Table S3. Source of discrepancy between datasets**

The outlying genes of the venn diagram excluding those unique to our TraDIS data. Gene names used by the Keio collection are shown in brackets. Venn subcategory is indicated in column 2 (K = Keio only; KP = Keio and PEC; KT = Keio and TraDIS (this study); P = PEC only; PT = PEC and TraDIS).

**Table S4. Essential genes identified by TraDIS analysis after outgrowth in LB.**

## Supplementary Methods

### Statistical Analysis

A Poissonian model, parameterized only by insertion density, gives the probability that no insertions will be found in a single region of a given length (for example, 60 bases), whereas genes and genomes have many such regions that are effectively independent. Briefly, if a p-value of 0.05 is associated with an IFR of length 60 under a simple Poisson model, every length-60 region in the gene or genome of interest has around a 5% chance of being insertion-free by chance under the null hypothesis. As a genome will contain many thousand such regions, many IFRs of length 60 will be expected by chance under the null hypothesis. A simulation of random insertion events reveals ~2900 IFRs of length 60 or above by random chance in a 4.8Mb genome with 370, 000 insertions (red markers in Fig. S1A).

To correct for this, we need to carefully state the statistic of interest and the corresponding null model. At least three probabilities are pertinent here: those with which, under a null model of random, independent insertions, (i) a single length-l region has no insertions; (ii) a gene of length g contains one or more IFRs of length l; (iii) a genome of length G contains one or more IFRs of length l. Previous models have implicitly estimated only (i), but to control the false discovery rate for individual genes or on a genome-wide basis, (ii) and (iii) respectively, are required. An analytic calculation of the corrected p-values corresponding to (ii) and (iii), which we label p_gene and p_genome, is laborious; for simplicity and illustrative power we computationally investigate these statistics, simulating many instances of N random non-coincident insertions in a genome of length G and reporting the statistics of resultant IFRs (C code available as SI). Reinterpreting the statistics in previous papers on this topic, and assuming a representative gene length of g = 1000, we find (Fig. S1) that in Langridge *et al*. (371,775 inserts, 4,791,961 base genome), the null model of genome-wide random insertion gives an expected ~16,000 IFRs with l >= 39 (classified in the original study using the Poisson model as p < 0.05) and ~2,900 IFRs with l >= 60 (classified as p < 0.01); the corrected lengths for p_genome = 0.05 and p_gene = 0.05 are ~200 and ~85 respectively (Figures S1B,D; [1]). In Barquist *et al*. (549,086 inserts, 4,878,012 base genome), the null model of genome-wide random insertion gives an expected ~22,000 IFRs of l >= 27 (classified as p < 0.05) and ~4,100 IFRs of l >= 41 (classified as p < 0.01); the corrected lengths for p_genome = 0.05 and p_gene = 0.05 are ~135 and ~62 respectively [2]. Our study (901,383 inserts, 4,631,469 base genome) gives a corrected p_genome = 0.05 of ~75 (and p_gene = 0.05 of ~36).

A further complication arises because the probability of observing an IFR of given length in a gene and the labelling of that gene as essential are not trivially related. Rather, genes are classified as essential if their insertion score clusters in the low-score category. While the analysis above can therefore give an indication of the resolution of our experiment, further theory, perhaps grounded in Hidden Markov Model analysis [3], will be required to forge a rigorous connection between the statistics of IFRs and the statistical power associated with classification of genes.

